# An image processing pipeline for electron cryo-tomography in RELION-5

**DOI:** 10.1101/2024.04.26.591129

**Authors:** Alister Burt, Bogdan Toader, Rangana Warshamanage, Andriko von Kügelgen, Euan Pyle, Jasenko Zivanov, Dari Kimanius, Tanmay A.M. Bharat, Sjors H.W. Scheres

## Abstract

Electron tomography of frozen, hydrated samples allows structure determination of macromolecular complexes that are embedded in complex environments. Provided that the target complexes may be localised in noisy, three-dimensional tomographic reconstructions, averaging images of multiple instances of these molecules can lead to structures with sufficient resolution for *de novo* atomic modelling. Although many research groups have contributed image processing tools for these tasks, a lack of standardisation and inter-operability represents a barrier for newcomers to the field. Here, we present an image processing pipeline for electron tomography data in RELION-5, with functionality ranging from the import of unprocessed movies to the automated building of atomic models in the final maps. Our explicit definition of metadata items that describe the steps of our pipeline has been designed for inter-operability with other software tools and provides a framework for further standardisation.

## Introduction

In the electron cryo-tomography (cryo-ET) approach, a three-dimensional (3D) reconstruction called a tomogram, is calculated from a series of images that are taken as a sample is rotated around a tilt axis in the electron microscope. Because radiation damage limits the dose that can be applied to the specimen, tomograms typically suffer from large amounts of noise. Moreover, samples are typically thin, slab-like sections, which dictates that some views of the sample cannot be acquired, leading to artefacts in the reconstructed tomogram. Provided that the structures of interest, or particles, can be localised in the noisy tomograms and that they can be brought into register, many instances of a particle can be combined into a single 3D reconstruction with an increased signal to noise ratio. This approach, commonly referred to as subtomogram averaging, has been used to calculate structures of macromolecular complexes to resolutions sufficient for de novo atomic modelling for macromolecules inside cells ^1–3^.

Many tools exist for acquiring and analysing cryo-ET data. SerialEM ^4,5^, one of the most popular softwares for tilt series acquisition, saves acquisition-related metadata in text files with the mdoc extension. Tomo5, a commercial software provided by Thermo Fisher, also writes metadata in the mdoc format. Accurate 3D reconstruction from a tilt series requires determining the parameters of a projection model describing the position, orientation and possibly also deformation of the sample in the electron microscope. The IMOD software package ^6,7^ provides some of the most popular tools for this task, which is commonly referred to as tilt series alignment. Recently, AreTomo ^8^ has also gained popularity by providing robust and automated tools for tilt series alignment, particularly for *in situ* data without fiducial markers. Once the parameters of a projection model are determined, a tomogram may be reconstructed from tilt images. Again, many approaches exist, including weighted back-projection ^9^, simultaneous iterative reconstruction technique ^10–12^, algebraic reconstruction technique ^13–15^ and compressed sensing ^16^. The reconstruction method used depends on what the tomogram, or sub-volumes thereof, will be used for. For example, tomograms with high contrast at lower spatial frequencies are often used for direct visual interpretation and segmentation, whilst for subtomogram averaging, high-resolution features need to be preserved in the reconstruction and lower resolution features may be less pronounced. Direct reconstruction of local regions of interest in tomograms ^1,17–20^ has obviated the need for calculating large, high-resolution tomograms from which subvolumes are cropped. To assist in feature identification, deep-learning approaches have gained traction for denoising ^21–24^ and template matching ^25,26^.

Despite the advent of near-complete cryo-ET image processing workflows, like those implemented in IMOD ^7^, PEET ^27^, EMAN2 ^28^, Dynamo ^29^, STOPGAP ^30^, TomoBEAR ^31^, SCIPION ^32,33^ or NextPyp ^19^, a lack of standardisation represents a hurdle to newcomers and makes it difficult to effectively use the best parts of each package.

Here, we present a cryo-ET image processing pipeline inside the free, open-source software RELION-5. Closely mirroring similar procedures in single-particle analysis, our cryo-ET pipeline starts from unprocessed movies and acquisition metadata in mdoc files and ends with high-resolution 3D reconstructions and the automated building of atomic models ^34^. The pipeline provides wrappers to CTFFIND4 ^35^ for estimation of contrast transfer function (CTF) parameters; to IMOD ^7^ and AreTomo ^8^ for tilt series alignment; and to cryoCARE ^22^ for tomogram denoising. We present graphical tools for the manual curation of images in tilt series and for the picking of particles in reconstructed tomograms. New to RELION-5 is the option to write out CTF-pre-multiplied 2D stacks of sub-images that are cropped from the tilt series at the corresponding positions for each particle. 2D particle stacks can then be used in the same alignment and classification procedures that were introduced in RELION-4 ^18^. Refinement of 2D particle stacks provides major computational advantages, both in disk space requirements and in processing speed, compared to methods that reconstruct 3D subvolumes, such as the pseudo-subtomogram approach introduced in RELION-4 ^18^. We also provide an explicit definition of the metadata required for and generated in our pipeline, providing a framework for further standardisation that will also be adopted in the CCP-EM software ^36,37^ (Tom Burnley, personal communication).

## Methods

All new developments in the RELION-5 tomography pipeline are accessible from the graphical user interface (GUI; Figure 1), which can be launched from the command line using the command `‘relion –tom’`. The side bar in the top left of the GUI shows the names of all job types. Below, we describe all jobs that have new developments for cryo-ET in RELION-5.

**Figure 1:**
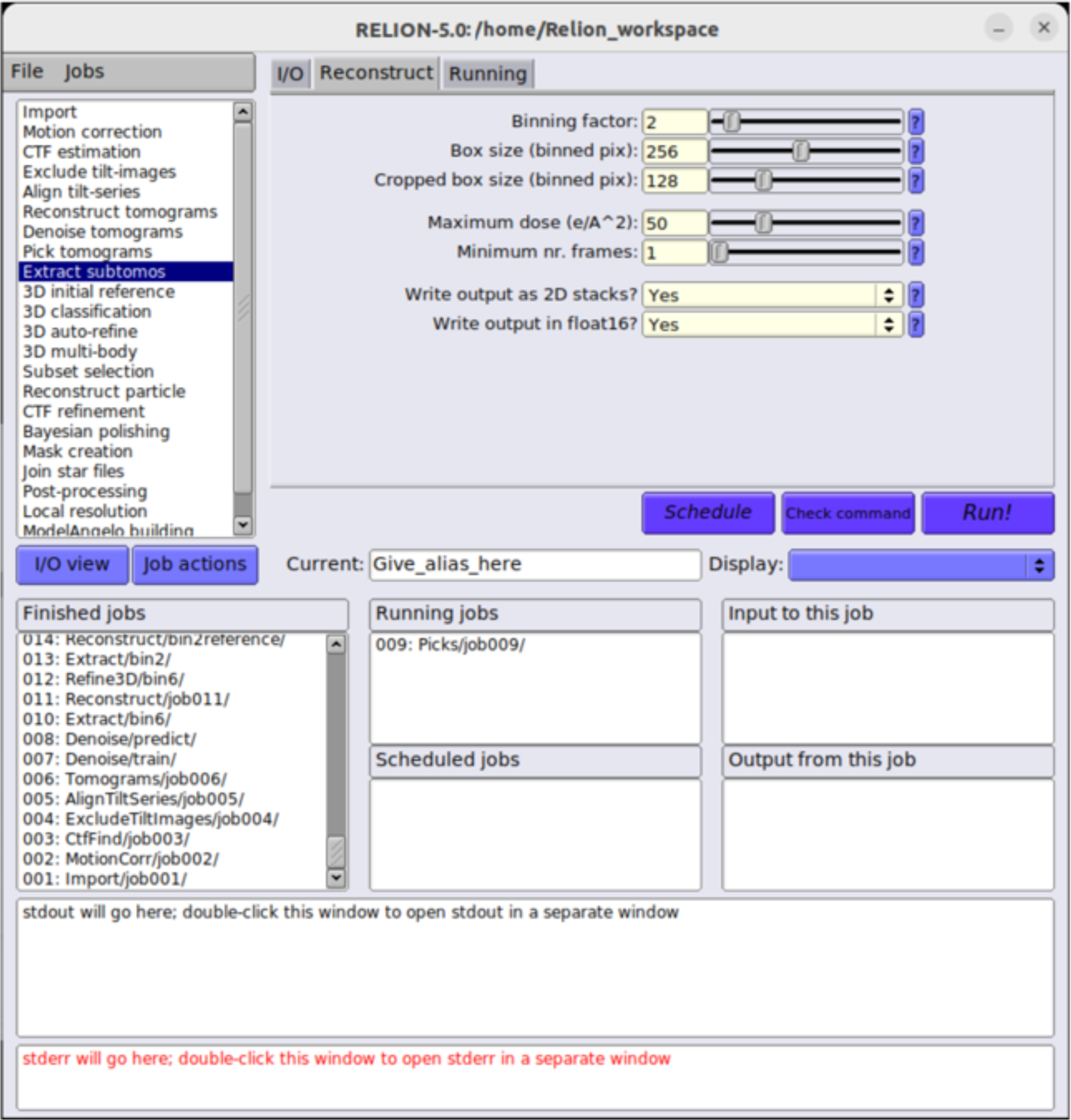
The RELION-5 tomography GUI. The sidebar on the top left provides access to all job types. The top right panel provides multiple tabs with input parameters for each job type. The bottom half of the GUI provides an overview of finished, running and scheduled jobs; how different jobs relate to each other through their input and output; and the output from the currently selected job.

### Importing into RELION’s data model

Starting from directories with the unprocessed movie files and their corresponding mdoc files, the “Import” job in the cryo-ET pipeline of RELION-5 writes metadata files that describe the acquired tilt series in the STAR format ^38^. A primary `tilt_series.star` file (Figure 2; Table 1) contains a global table with one line per tilt series. The global table provides general information, including the acceleration voltage, pixel size, amplitude contrast and spherical aberration, plus a reference to a STAR file that contains information about individual images in each tilt series (Figure 3; Table 2). Tilt series STAR files are stored in the `tilt_series/` subdirectory of the Import job directory. They contain a table with one line for each image of the tilt series that stores the name of the unprocessed movie, the nominal tilt angle of the stage, the nominal orientation of the tilt axis, the nominal defocus and the accumulated electron dose (Figure 3; black).

**Figure 2:**
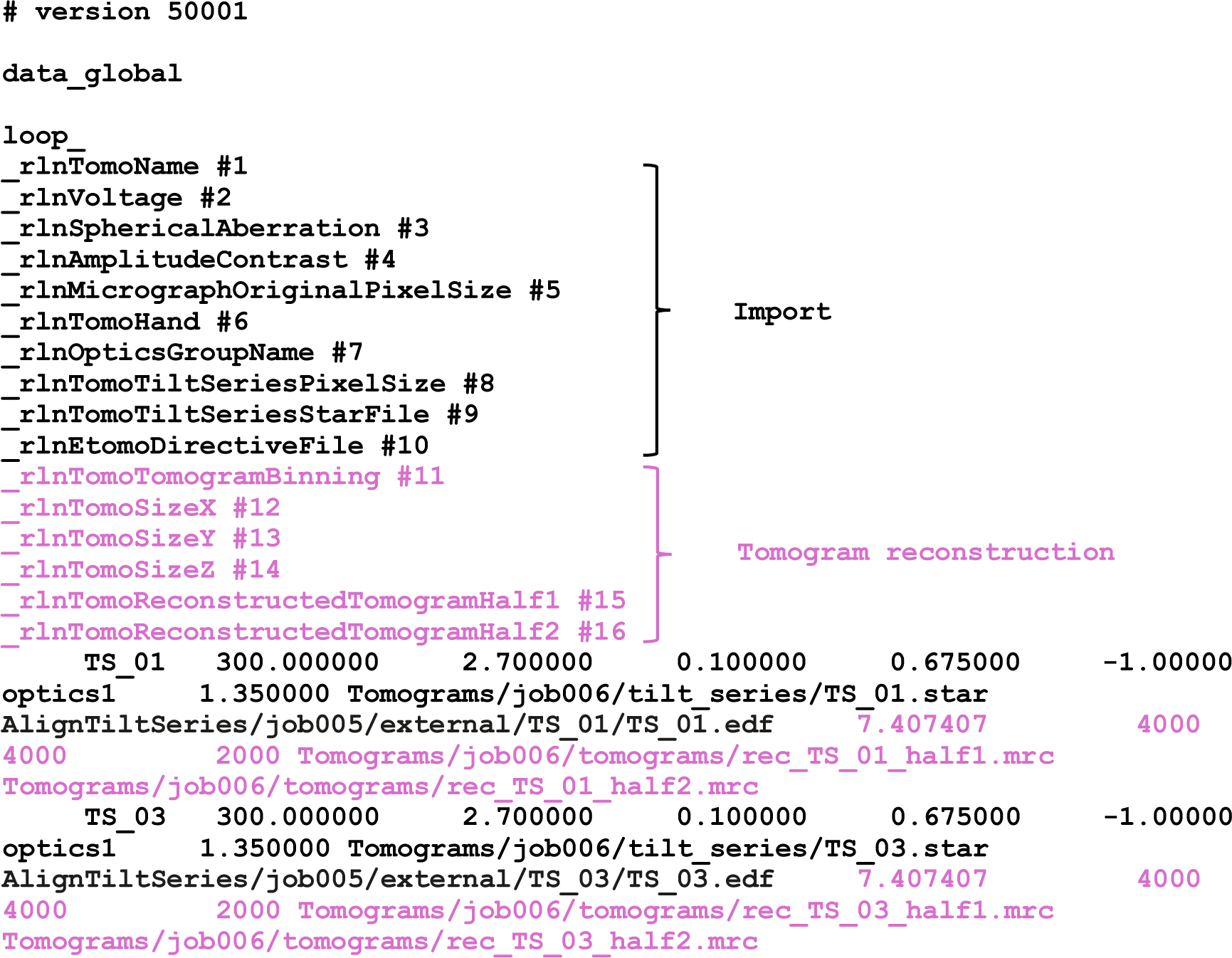
Tomograms metadata description. Metadata to describe a set of tomograms is stored in the STAR format. Labels and data columns shown in black are added during import; labels and columns in violet are added during tomogram reconstruction. Corresponding definitions of metadata labels are given in Table 1.

**Figure 3:**
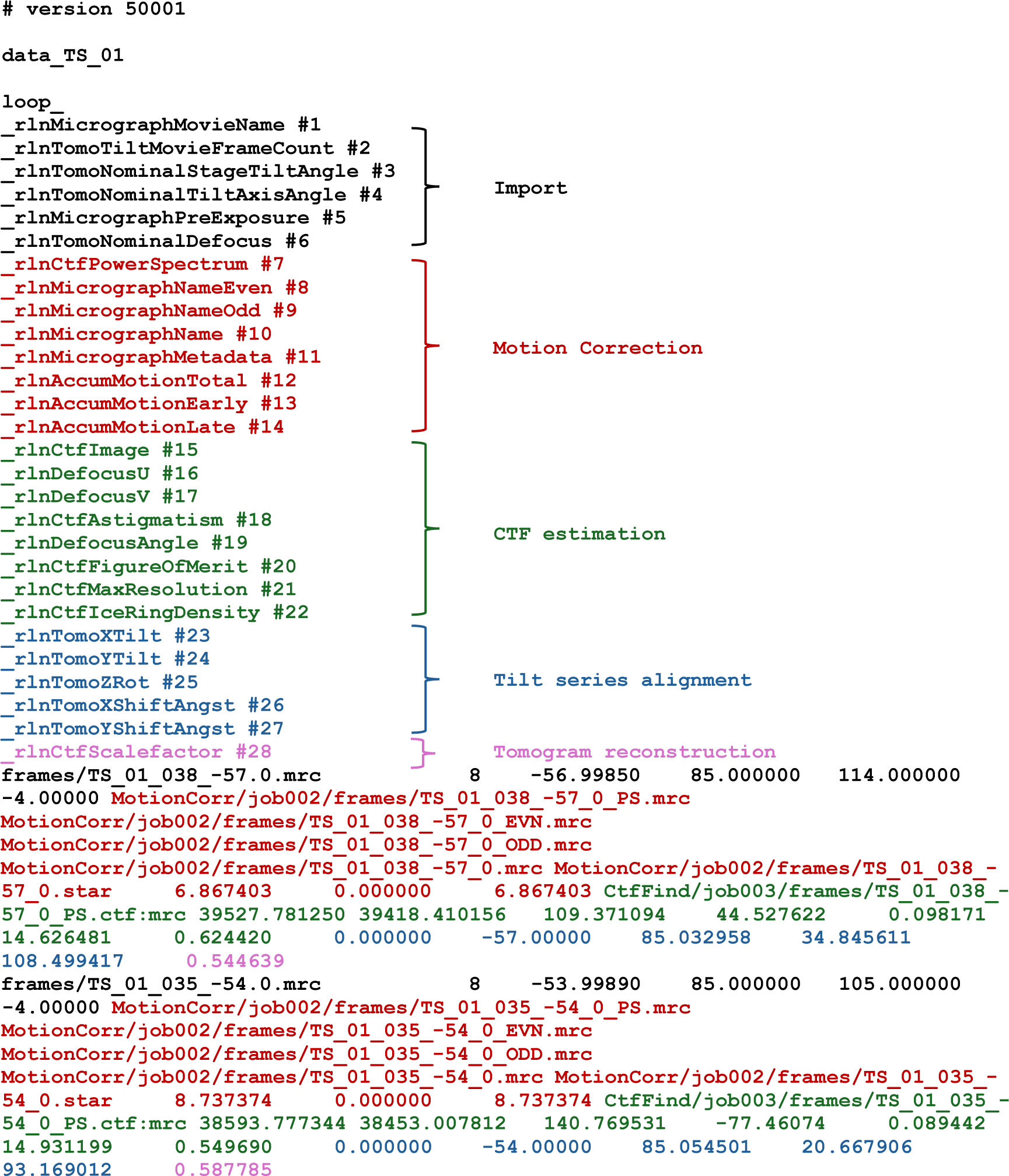
Tilt series images metadata description. Metadata to describe a set of images in a tilt series is stored in the STAR format. Labels and data columns shown in black are added during import; labels and columns in red are added during motion correction; labels and columns in green are added during CTF estimation; labels and columns in blue are added during tilt series alignment; labels and columns in violet are added during tomogram reconstruction. Corresponding definitions of metadata labels are given in Table 2.

**Table 1:**
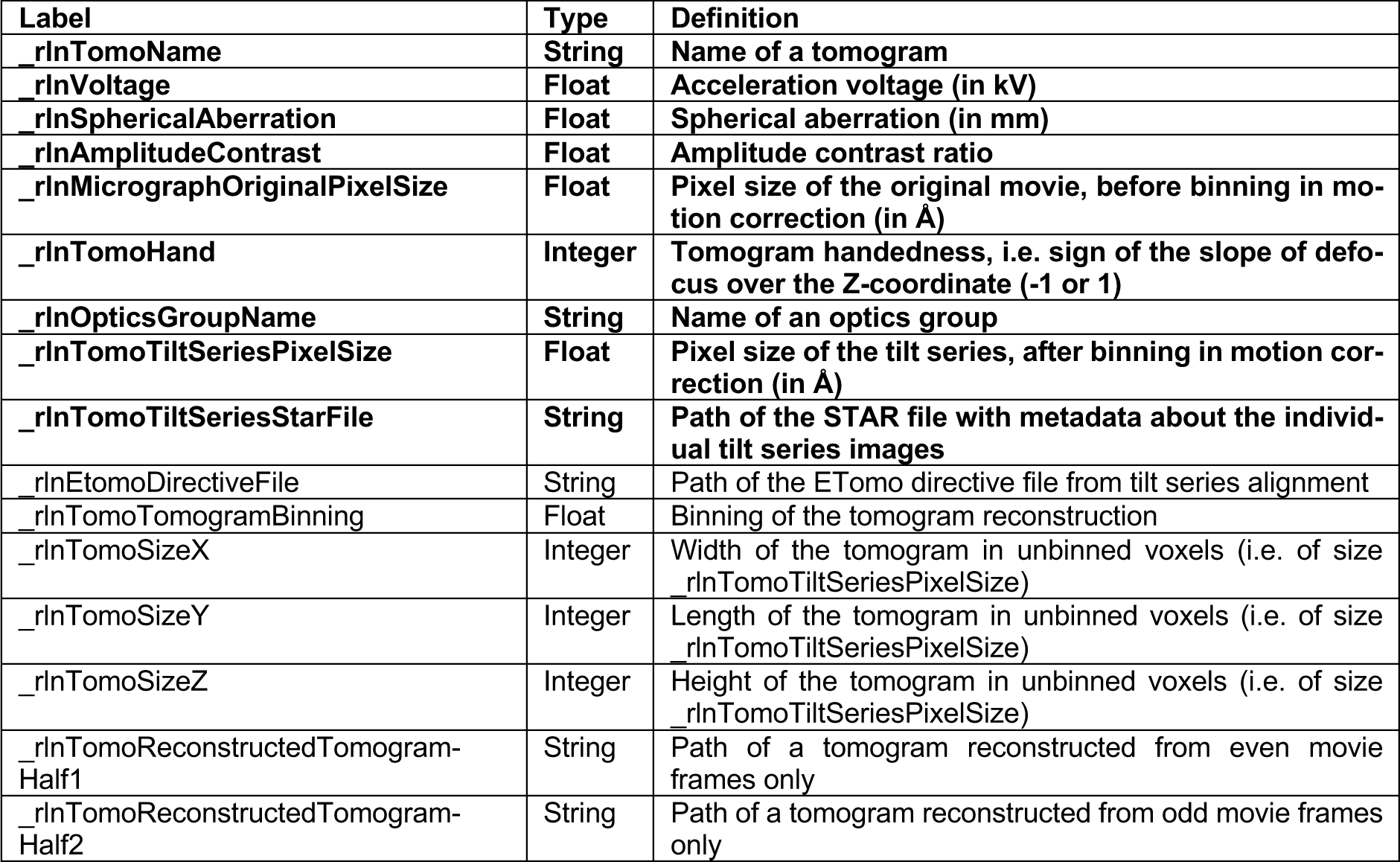
Metadata items for tomograms. Metadata labels, types and definitions for the description of a set of tomograms. Entries in bold are compulsory for sub-tomogram averaging; entries in regular are not necessary for sub-tomogram averaging.

**Table 2:** Metadata items for tilt series images. Metadata labels, types and definitions for the description of a set of images in a tilt series. Entries in bold are compulsory for sub-tomogram averaging; entries in regular are not necessary for sub-tomogram averaging.

| Label | Type | Definition |
| --- | --- | --- |
| `_rlnMicrographMovieName` | String | Path of the movie stack for an individual tilt series image |
| `_rlnTomoTiltMovieFrameCount` | Integer | Number of frames in the movie stack |
| `_rlnTomoNominalStageTiltAngle` | Float | Nominal value for the stage tilt angle (in °) |
| `_rlnTomoNominalTiltAxisAngle` | Float | Nominal value for the angle of the tilt axis with the Y-axis (in °) |
| **`_rlnMicrographPreExposure`** | **Float** | **Pre-exposure dose (in electrons/Å<sup>2</sup>)** |
| `_rlnTomoNominalDefocus` | Float | Nominal value for the defocus (in µm, negative values for underfocus) |
| `_rlnCtfPowerSpectrum` | String | Path of the power spectrum image for CTF estimation |
| `_rlnMicrographNameEven` | String | Path of the summed micrograph image from even movie frames only |
| `_rlnMicrographNameOdd` | String | Path of the summed micrograph image from odd movie frames only |
| **`_rlnMicrographName`** | **String** | **Path of the summed micrograph image from all movie frames** |
| `_rlnMicrographMetadata` | String | Path of a STAR file with metadata from motion correction |
| `_rlnAccumMotionTotal` | Float | Accumulated global motion during the entire movie (in Å) |
| `_rlnAccumMotionEarly` | Float | Accumulated global motion during early frames of the movie (in Å) |
| `_rlnAccumMotionLate` | Float | Accumulated global motion during late frames of the movie (in Å) |
| `_rlnCtfImage` | String | Path of the Thon-ring image from CTF estimation |
| **`_rlnDefocusU`** | **Float** | **Defocus in the U-direction (in Å, positive values for underfocus)** |
| **`_rlnDefocusV`** | **Float** | **Defocus in the V-direction (in Å, positive values for underfocus)** |
| `_rlnCtfAstigmatism` | Float | Absolute value of the difference between defocus in U- and V-direction (in Å) |
| **`_rlnDefocusAngle`** | **Float** | **Angle between X and defocus U direction (in °)** |
| `_rlnCtfFigureOfMerit` | Float | Figure of merit from CTF estimation |
| `_rlnCtfMaxResolution` | Float | Maximum resolution (in Å) of fitted Thon rings from CTF estimation |
| `_rlnCtfIceRingDensity` | Float | Accumulated power of the image in the frequency range (0.25-0.28 Å<sup>-1</sup>) |
| `_rlnTomoXTilt` | Float | Angle for rotation of the tomogram around the X-axis (in °) |
| `_rlnTomoYTilt` | Float | Angle for rotation of the tomogram around the Y-axis (in °) |
| `_rlnTomoZRot` | Float | Angle for rotation of the tomogram around the Z-axis (in °) |
| `_rlnTomoXShiftAngst` | Float | Shift in X-direction (in Å) to align the projection of a tomogram with the tilt series image |
| `_rlnTomoYShiftAngst` | Float | Shift in Y-direction (in Å) to align the projection of a tomogram with the tilt series image |
| `_rlnCtfScalefactor` | Float | Linear scale-factor to be applied to the CTF values |

One option that the user needs to decide when importing tilt series is the defocus handedness (`rlnTomoHand` in Figure 2; Table 1), which defines the direction in which defocus changes moving across the tilt axis in tilted images. Although this could be estimated automatically during CTF estimation ^39^, for now, the defocus handedness needs to be determined by the user using trial and error. For all Thermo Fisher Krios and Glacios microscopes that we have tested so far, the `rlnTomoHand` was always −1, which is the default option on the GUI (Yes, to invert defocus handedness).

### Motion correction

The `tilt_series.star` file from the “Import” job can be used as input for a “Motion correction” job. Internally, all images of all tilt series are combined into a single list of images that can then be processed in parallel using a combination of message passing interface (MPI) and threads. Either UCSF MotionCor2 ^40^ or RELION’s own implementation of that algorithm ^41^ may be used for movie frame alignment. An option exists to write out summed micrographs that are calculated only from the even or the odd movie frames, which may be used for tomogram denoising as described below. Although an option exists to bin images at this stage, this option will limit the achievable resolution of all downstream steps and is therefore only used in case images were recorded in super-resolution mode.

The motion correction job adds several columns to the tables in the STAR files of the individual tilt series, including the name of the motion-corrected micrograph and statistics about the estimated movements in each of the tilt series images (Figure 3; red).

### CTF estimation

The `tilt_series.star` file from the “Motion correction” job can be used as input for a “CTF estimation” job, where individual images of all tilt series are again processed in parallel using MPI. CTF estimation is performed using a wrapper to CTFFIND4 ^35^, with two modifications from the equivalent approach for single-particle analysis that increase robustness for images at high tilt angles, where the signal to noise ratio is low. Firstly, whereas the same minimum and maximum defocus values are used for all micrographs in single-particle analysis, in the tomography pipeline the user may provide a search range that will be used around the nominal defocus value for each image. Secondly, the maximum resolution used for CTF estimation may be varied with the accumulated electron dose for each image in the tilt series.

This job again adds information to the STAR files of the individual tilt series, including the estimated defocus values, a figure-of-merit for the CTF fit, and the maximum resolution to which a good fit was obtained for each image of the tilt series (Figure 3; green). The values for `rlnCtfIceRingDensity` are calculated as the power of the tilt series images between 0.25 and 0.28 Å^−1^ ^42^.

### Selection of tilt series images

Given a `tilt_series.star` file, the “Exclude tilt-images” job launches a Napari-based ^43,44^ viewer that displays the images in a tilt series and allows the user to de-select unsuitable images, for example images in which the field of view has shifted or is obstructed by the grid bars, or if there is significant radiation induced movement detected. A screenshot of the viewer is shown in Figure 4a.

**Figure 4:**
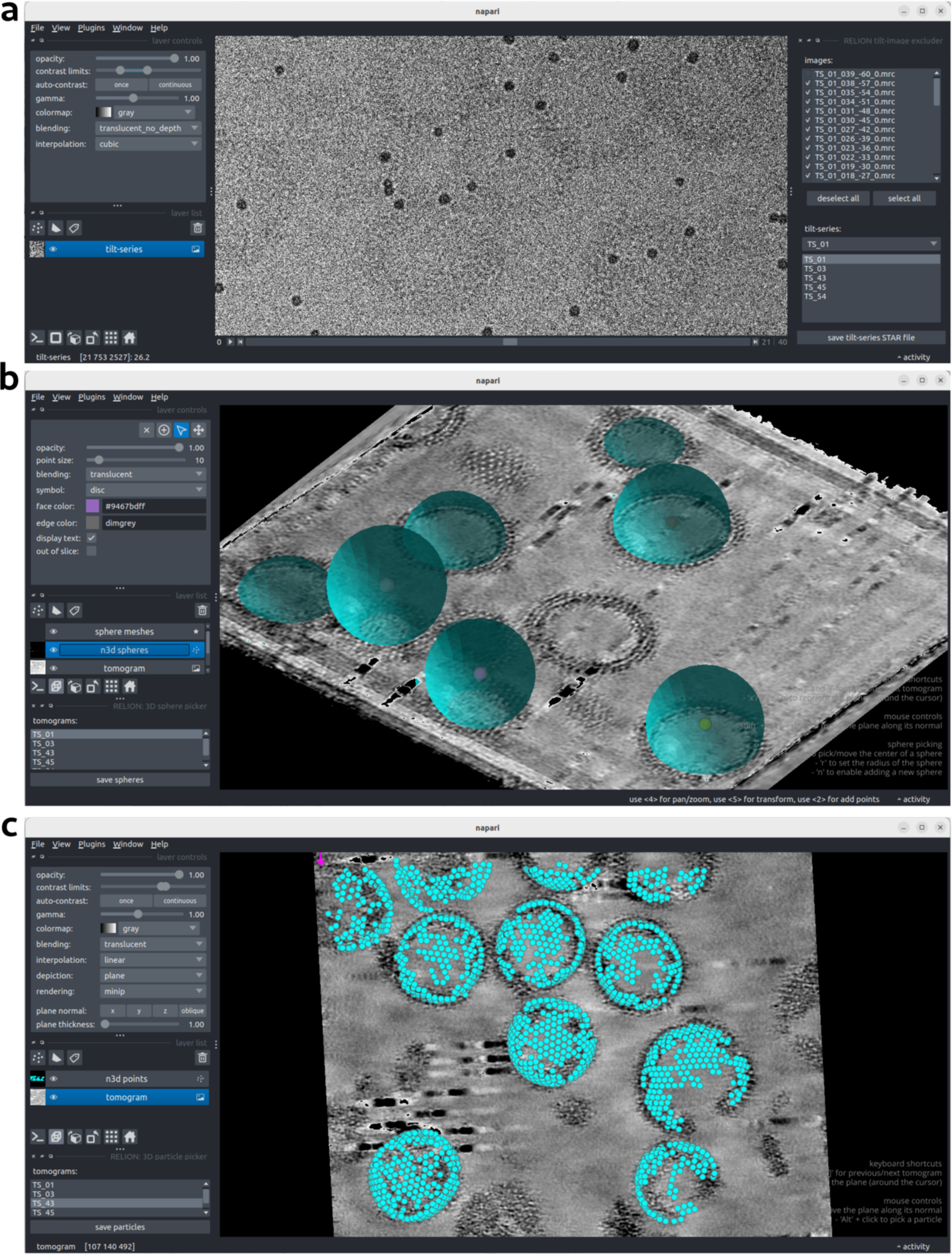
Screenshots of Napari-based picking tools. **(a)** The tilt-series selection program opened by the “Exclude tilt-images” job allows the user to de-select tilt images for each tilt series. **(b)** The picker plug-in opened by the “Pick tomograms” job allows one to annotate the reconstructed tomograms with spheres (shown here) or 1D-curves (not shown), which will then be used to randomly sample particles with priors that orient the normal to the sphere or along the curves with the Z-axis. **(c)** Individual particles can also be annotated manually, or read from a STAR file for visualisation.

The output selected_tilt_series.star file will point towards individual tilt series STAR files that no longer contain lines with de-selected images.

### Tilt series alignment

The “Align tilt series” job implements wrappers for performing tilt series alignment in IMOD ^7^ and AreTomo ^8^. In IMOD, fiducial markers or local image patches are tracked through a tilt series yielding observed 2D positions of a 3D object, and parameters of a projection model are fit to these observations. In AreTomo, a projection matching routine is used to iteratively improve projection model parameters. Each wrapper takes the information from an input `tilt_series.star` file and prepares the appropriate commands to align tilt series. This wrapper has not been parallelised.

The RELION-5 projection model at this stage is defined as 5 parameters per tilt image. `rlnTomoXTilt, rlnTomoYTilt` and `rlnTomoZRot` constitute a set of extrinsic Euler angles (in °), which rotate the specimen within a fixed microscope coordinate system. In this coordinate system, the optical axis is aligned with the Z-axis and the stage tilt axis is aligned with the Y-axis. The centre of rotation is the centre of the tomogram. The first rotation around the X-axis (`rlnTomoXTilt`) accounts for non-perpendicularity of the stage tilt axis to the optical (Z) axis. In most cases, this angle will be close to 0°. The second rotation around the Y-axis (`rlnTomoYTilt`) is the stage tilt angle. The third rotation around the Z-axis (`rlnTomo-ZRot`) aligns the Y axis to the tilt axis in the projection image. `rlnTomoXShiftAngst` and `rlnTomoYShiftAngst` are shifts (in Å from the centre of rotation) applied after rotating the specimen to align its projection to the experimental image data.

Denoting rlnTomoXTilt, rlnTomoYTilt and rlnTomoZRot as *ε_x_*, *ε_y_* and *ε_z_*, and rlnTomoXShiftAngst and rlnTomoYShiftAngst as *ι1_x_* and *ι1_y_*, respectively, the transformation matrix *R* that converts (centred) coordinates in the 3D tomogram to (centred) coordinates in the 2D tilt series images is thereby defined as:

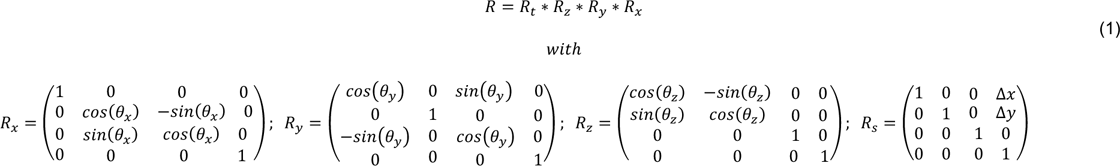

Alignment parameters are extracted from output files of IMOD or AreTomo, and written in an output aligned_tilt_series.star file (Figure 3; blue).

Python application programming interfaces for tilt series alignment procedures in RELION are available as standalone python packages yet-another-imod-wrapper (https://github.com/teamtomo/yet-another-imod-wrapper) and lil-aretomo (https://github.com/teamtomo/lil-aretomo).

### Tomogram reconstruction

The “Reconstruct tomograms” job takes an `aligned_tilt_series.star` file as input and uses real-space weighted back-projection, with pre-multiplication of the CTF using a single defocus value for each tilt image, to reconstruct the corresponding tomograms. As such, these tomograms will have small errors in defocus, which become worse as one moves away from the centre of the reconstruction. In the presented workflow, the reconstructed tomograms are only used for picking particles and subsequent averaging approaches use cropped 2D stacks as individual particles. Tomograms are therefore typically reconstructed with relatively large pixel sizes (e.g. 10 Ångstrom). If even/odd micrographs were calculated in the “Motion correction” job, then even/odd tomograms can also be calculated for denoising in the “Denoise tomograms” job below. In addition, an option exists to apply an overall tilt angle offset to the tilt series, which may be useful, for example, when tilt series were collected on lamellae that were milled from a thick specimen at a pre-selected angle away from the direction of electron beam using a focused ion beam (FIB).

The output `tomograms.star` file, besides containing all the information of the input tilt series file, contains links to the names of the reconstructed tomograms, which are saved in MRC format in a sub-directory of the job called `tomograms/`. This job also adds a column with a CTF scale-factor, calculated as the cosine of the tilt angle, to the STAR files of the individual tilt series images (Figure 3; violet).

### Denoising tomograms

The “Denoise tomograms” job takes a `tomograms.star` file as input and implements a wrapper to the noise-to-noise denoising program cryo-CARE ^45^. In the first part of the job, a neural network is trained on a selected subset of representative even/odd tomograms that were constructed in the “Reconstruct tomograms” job. In the second part of the job, the trained network is applied to the complete set of tomograms that were calculated from all movie frames. The output tomograms.star file from this job points to the denoised tomograms, which are again stored in a sub-directory of the job called `tomograms/`.

### Picking particles

A `tomograms.star` file is used as input for the “Pick tomograms” job, which Napari ^43^ as image viewer and napari-threedee ^46^ to provide tools for the interactive 3D annotation of isolated particles, filaments or spheres on tomogram slices, similar to dtmslice from Dynamo ^47^. Particles can be taken directly from particle annotations, or sampled from filaments or spheres. Picking on spheres (Figure 4b) is useful for particles that are arranged, for example, on spherical virus capsids, vesicles or cells. Filaments are picked as 1D curves. An option to sample particles from non-spherical 2D surfaces was planned, but has not yet been implemented. The napari viewer can also be used to visualise sets of particle coordinates after alignment and/or classification in the latter parts of the pipeline (Figure 4c). Sampling from different geometries is implemented in a standalone python package `morphosamplers` (https://github.com/mor-phometrics/morphosamplers).

The coordinates of manually picked particles, spheres or filaments of individual tomograms are saved in STAR files in a sub-directory called `annotations/`. When using spheres or filaments, particles are sampled from the spheres or filaments according to a user-specified inter-particle distance. The output `particles.star` file contains the 3D coordinates of all picked particles in all tomograms, with coordinates defined in Ångstroms relative to the centre of the tomogram (Figure 5; Table 3). When particles are sampled from spheres or filaments, the output STAR file will also contain Euler angles that define a rotation that orients the Z-axis of the particle parallel to the surface of the sphere or perpendicular to the long filament axis. Thereby, in subsequent particle alignment and classification, a 90° prior on the tilt angle will orient particles with their Z-axis normal to the sphere surface or parallel to the long filament axis. Having non-zero tilt angles prevents gimbal locks in the Euler angles during refinement in RELION.

**Figure 5:**
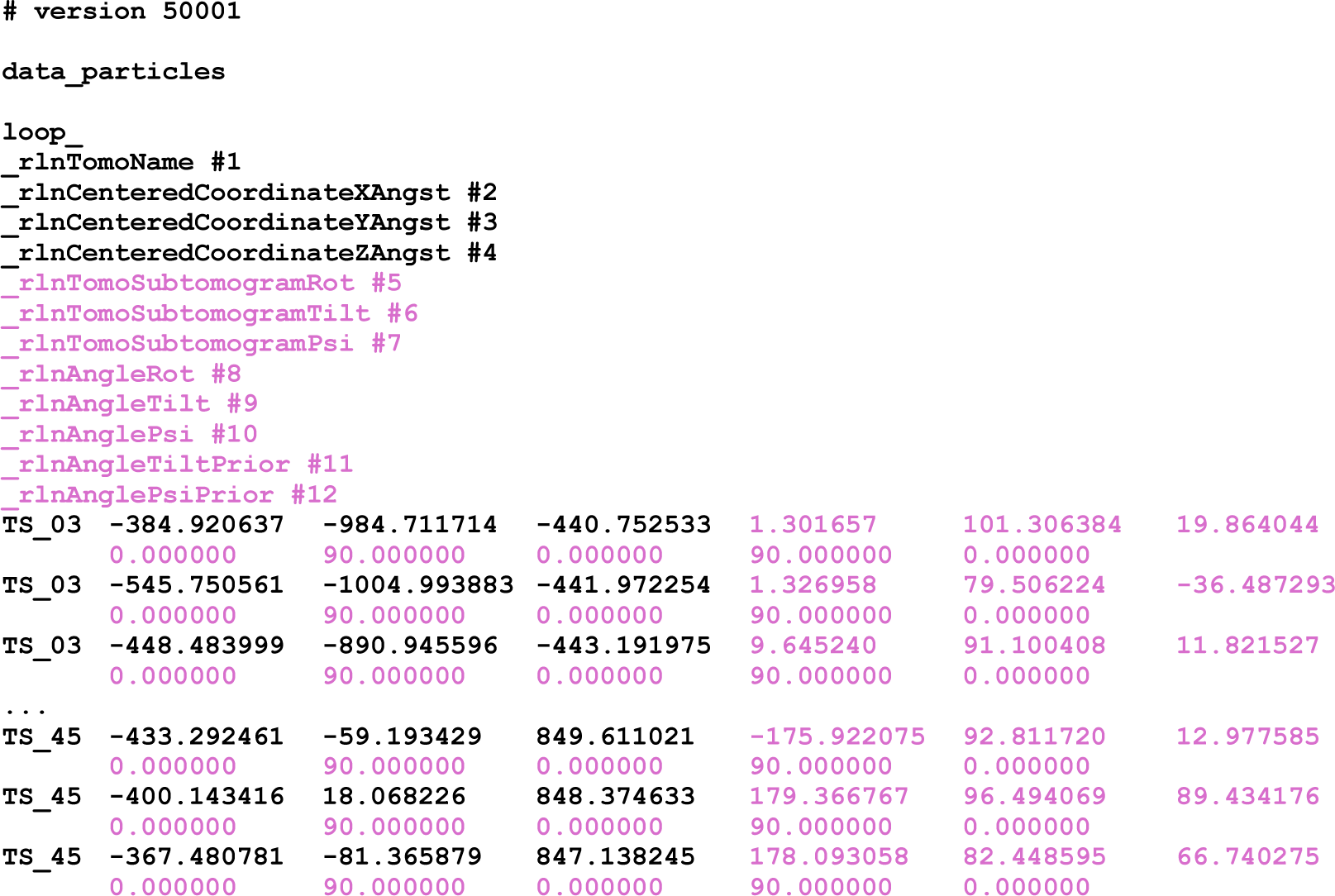
Particle metadata description after picking. Metadata to describe a set of particles after picking in the Napari GUI. A single STAR file contains particle coordinates (in Ångstroms relative to the centre of the tomogram) for all tomograms. The labels and data columns in violet arise from the sampling of the particles on manually picked spheres, and express information that aligns their Z-axis with the normal to the surface sphere. Corresponding definitions of metadata labels are given in Table 3.

**Table 3:**
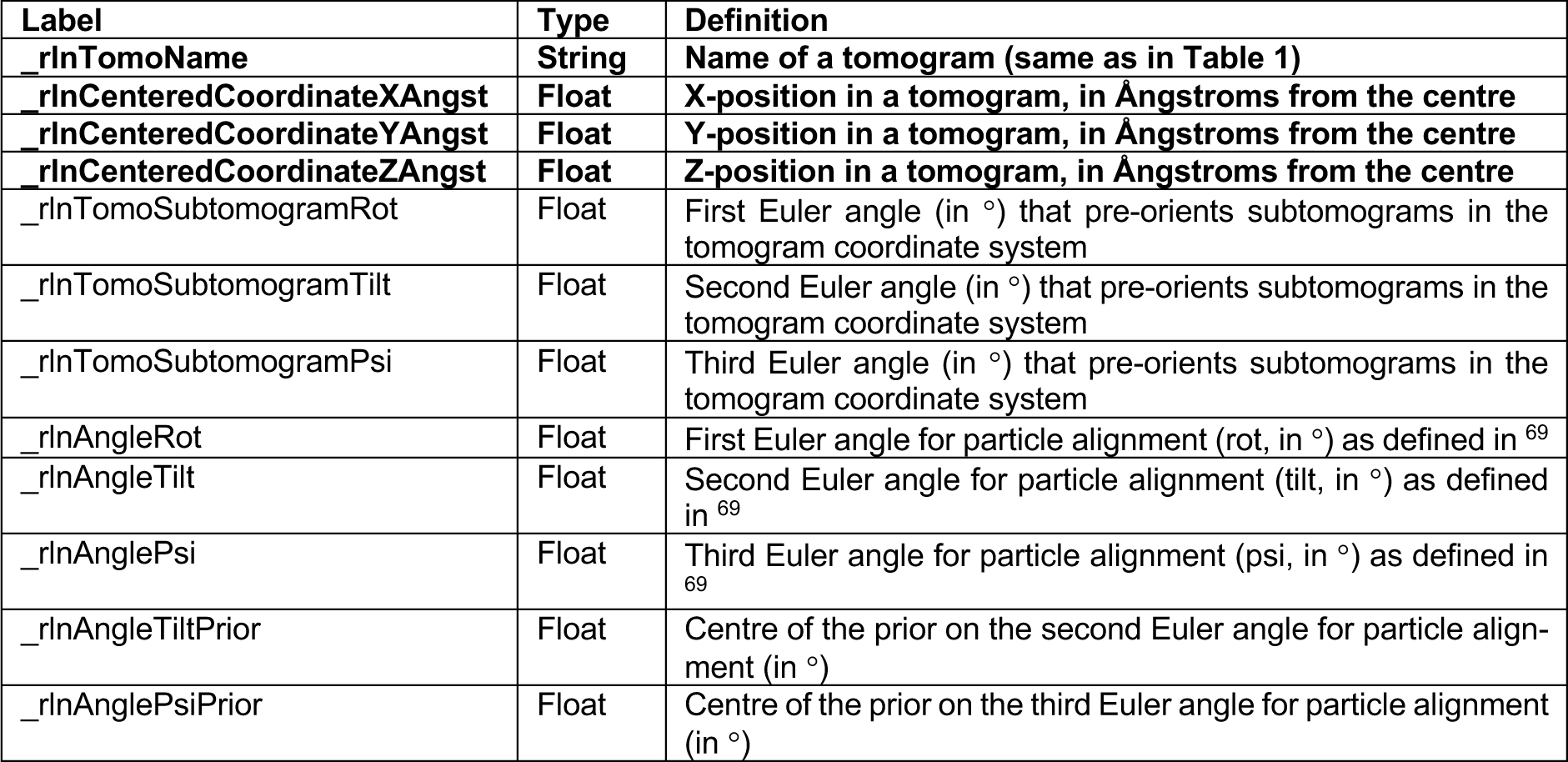
Metadata items for particle picks. Metadata labels, types and definitions for the description of a set of picked particles. Entries in bold are always present; entries in regular are only present when particles are sampled from spheres or filaments.

### Pseudo-subtomograms and 2D particle stacks

The `particles.star` file from the “Pick tomograms” job, together with the `tomograms.star` file from the “Reconstruct tomograms” job, form the input to the “Extract sub-tomos” job. Because these two files together define the input, they are bundled in a file called `optimisation_set.star`, which was also written out by the “Pick tomograms” job. The `optimisation_set.star` contains links to the filenames of both individual STAR files. One can either use the `optimisation_set.star` file or the two separate STAR files as input for the “Extract subtomos” job. This job will combine the 3D positions of the particles with the alignment parameters of the tomograms either to construct 3D pseudo-subtomograms (as introduced in RELION-4 ^18^), or to crop the relevant regions around individual particles in the tilt series images and save these as 2D particle stacks. Both the pseudo-subtomograms and the 2D stacks will be pre-multiplied with the CTF in images with a user-defined box size, and then possibly cropped to a smaller box size after the signal delocalisation has been compensated. These calculations are parallelised using both MPI and threads. Because there are typically fewer images in the tilt series (*N_tilt_*) than the cropped boxed size (*B*), 2D stacks (with *N_tilt_ * B*^2^ pixels) often occupy less disk space than 3D pseudo-subtomograms (with *B*^3^ voxels). Options exist to only use tilt series images below a user-defined maximum dose and to only output particles that are visible on a user-specified minimum number of tilt series images. Writing particles in float16 saves a factor of 2 in disk space compared to writing in the default float32 MRC format, although not all third-party programs may be able to read such images. The particle images are saved in a subdirectory of the job called `Subtomograms/`. The job also saves a `particles.star` file with the names of the particle images, and (for 2D stacks only) an array with names of images of the tilt series saved in the stacks. The job also writes out a new `optimisation_set.star` with the filenames of the output `particles.star` file and the input `tomograms.star` file.

### Subsequent averaging approaches

The `optimisation_set.star` from the “Extract subtomos” job can be used as input to any of the existing “3D initial reference”, “3D classification”, “3D auto-refine”, or “3D multi-body” jobs in the standard RELION pipeline. These methods already existed in single-particle and tomography pipelines of RELION ^18,41^ and are not described in detail here. Two new features are an option to resize input 3D reference maps and mask if they have a different pixel and/or box size than the input images (the same also works for standard single-particle analysis), and an option to impose a prior on the tilt angle, which is useful if particles were sampled from spheres or filaments. Each of these jobs will also output `optimisation_set.star` files with links to the STAR files with the tomograms and the refined particle coordinates.

### Bayesian polishing and CTF refinement

The methods for Bayesian polishing, which performs tilt series re-alignment and estimation of per-particle motion throughout the tilt series (but does not follow motion of individual particles through individual movie frames), and CTF refinement, which refines the defocus estimates for each tilt series image, have been described previously ^18^ and will not be discussed here. Besides an input `optimisation_set.star` file, these methods also need reference halfmaps, a reference mask and a `postprocess.star` file from a “Post-processing” job as input. Because these calculations are sensitive to the size and grey-scale of the reference, the reference maps need to be calculated by the “Reconstruct particle” job, and a binning factor of 1 must be used. A larger binning factor may be used in the preceding “3D classification” or “3D auto-refine” jobs.

The “CTF refinement” job will update the defocus values for all tilt series images and store links to updated STAR files for the individual tilt series in an updated `tomograms.star` file. Likewise, the “Bayesian polishing” job will update the tilt series alignment parameters in a new `tomograms.star` file. If per-particle motion estimation is performed, then the “Bayesian polishing” job will also output a `motion.star` file with the estimated per-particle motion tracks for all particles. A link to these files will also be added to the output `optimisation_set.star` file.

If one wishes to perform further refinements or classifications after a “CTF refinement” and/or a “Bayesian polishing” job, new particles should be extracted using the “Extract subtomos” job with the updated CTF parameters and/or tilt series alignment parameters and particle positions. If no further refinement or classification is deemed necessary, then calculating a new reconstruction with the “Reconstruct particle” job will suffice.

## Results

We illustrate the capability of our pipeline on a dataset of five tomograms of virus-like particles (VLPs) of the capsid and spacer peptide 1 (CA-SP1) region of the Gag polyprotein in the immature human immunodeficiency virus 1 (HIV-1) ^48^. The same subset of five tomograms has previously been used to benchmark various cryo-ET tools. NovaCTF introduced 3D CTF correction, leading to 3.9 Å resolution ^49^; optimizing frame alignment and CTF parameters in Warp led to 3.8 Å ^50^; a combination of Warp-RELION-M and Dynamo yielded 3.4 Å ^51^; and RELION-4 gave 3.2 Å ^18^.

Starting from the import of raw movie frames and SerialEM mdoc files, all processing steps were performed using the pipeline described in the **Methods** section. Movie frames were aligned using RELION’s own motion-correction program; tilt series images without signals (dark images) were manually excluded from the first two tilt series; CTF parameters were estimated using CTFFIND4 ^35^; and tilt alignment was performed using the fiducial-based procedure in IMOD ^7^. Next, the five tomograms were reconstructed with a binned voxel size of 10 Å and the napari-based 3D picker was used to manually annotate 47 spheres coinciding with the outer surfaces of the VLPs observed in the tomograms. Using an inter-particle distance of 60 Å, 30,597 particles were extracted by randomly sampling the spheres. Priors on the orientations of the particles, which position the normal to the sphere parallel to the Z-axis for each particle at a tilt angle of 90°, allowed us to obtain an initial 3D reference map by simply running a “Reconstruct particle” job with 6-fold rotational symmetry. The initial model was subjected to a first 3D auto-refine job at a binning factor of 6, followed by a second 3D auto-refine job at a binning factor of 2. After removing 8,937 duplicated particles (using the corresponding option in a Subset selection job), we then ran a 3D classification with 9 classes, local alignment and regularisation parameter T = 1. Manual selection of the best 2 classes resulted in a subset of 9,053 particles. These particles were used for a 3D auto-refinement at a binning factor of 1, which gave a map at 4.0 Å resolution. Five iterations of CTF refinement, Bayesian polishing and 3D auto-refinement at a binning factor of 1 yielded a final map at 3.3 Å. Automated model building with ModelAngelo^34^ led to an atomic model comprising 211 (94%) of the 224 residues that were built in the original study ^48^.

Fourier shell correlation (FSC) curves of some of the intermediate steps and the final reconstruction are shown in Figure 6a and the final map is shown in Figure 6b. In Figure 6c, we show representative regions from the reconstructed map after the first 3D refinement step at binning factor 1 (first row), after the first iteration of CTF refinement, Bayesian polishing and 3D auto-refinement (second row), and in the final map (third row), as well as the final map with the atomic model as output by the ModelAngelo job, with no additional model refinements. The computational costs of the individual steps in the pipeline are summarised in Table 4. For all 3D auto-refinements and the 3D classification leading to the final map, individual particles were written as 2D image stacks in the corresponding “Extract subtomos” jobs. For comparison with RELION-4, we also extracted particles at a binning factor of 1 as 3D pseudo-sub-tomograms. This required 20 times more disk space and resulted in an 8-fold increase in wall-clock compute time for 3D auto-refinement, compared to extracting particles as 2D image stacks.

**Figure 6:**
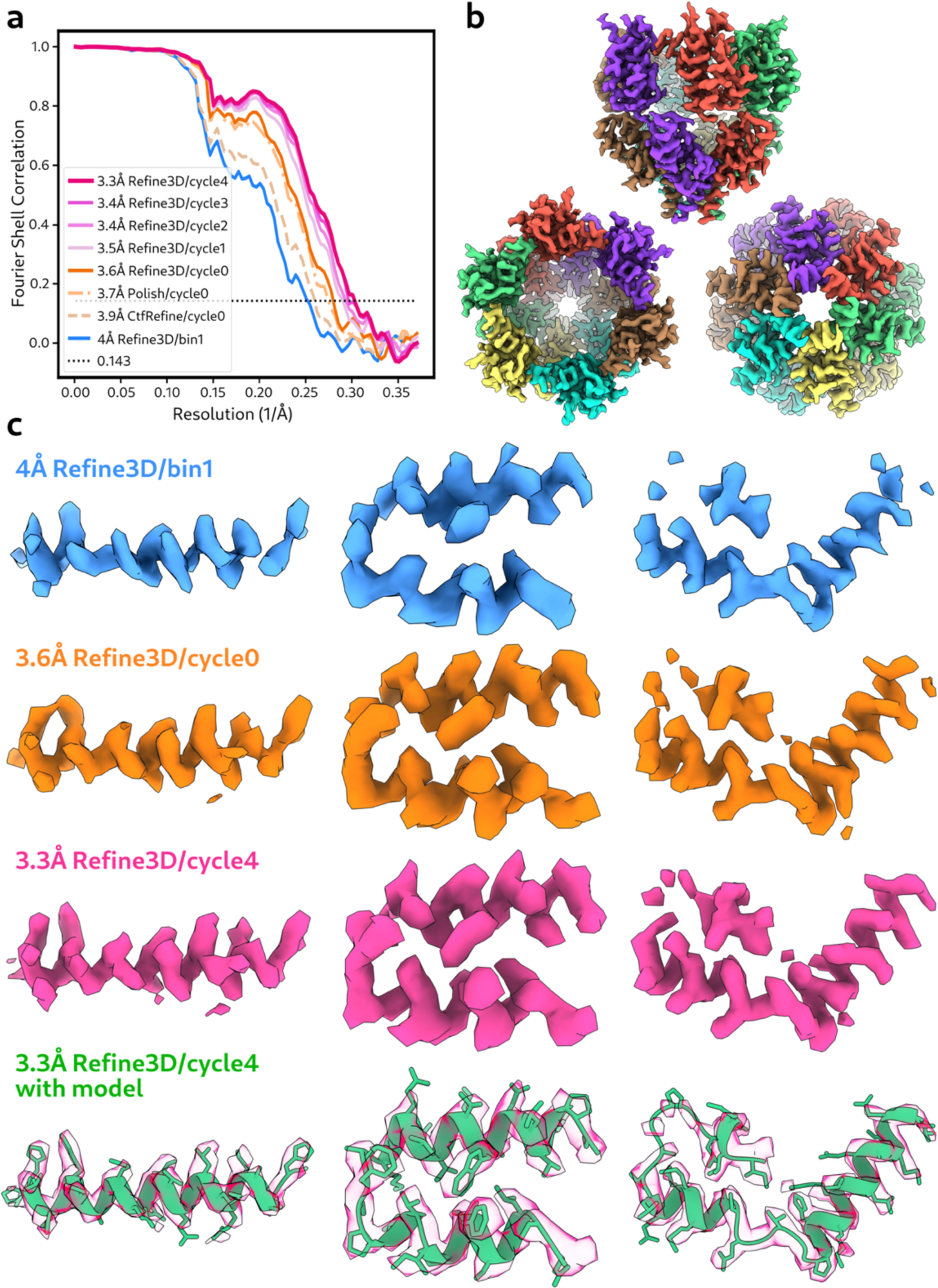
Subtomogram averaging of the CA-SP1 hexamer of the immature HIV1 capsid. **(a)** FSC curves at intermediate and final refinement steps in the RELION-5 pipeline. **(b)** Final reconstructed map, side view (top), top (bottom left) and bottom views (bottom right). **(c)** Representative regions in the reconstructed map after the first 3D auto-refinement at binning factor of 1 (blue, top row), after the first iteration of CTF refinement, Bayesian polishing and 3D auto-refinement (orange, second row), after the fifth iteration (magenta, third row), and the final map with the atomic model generated by ModelAngelo (green, bottom row).

**Table 4:**
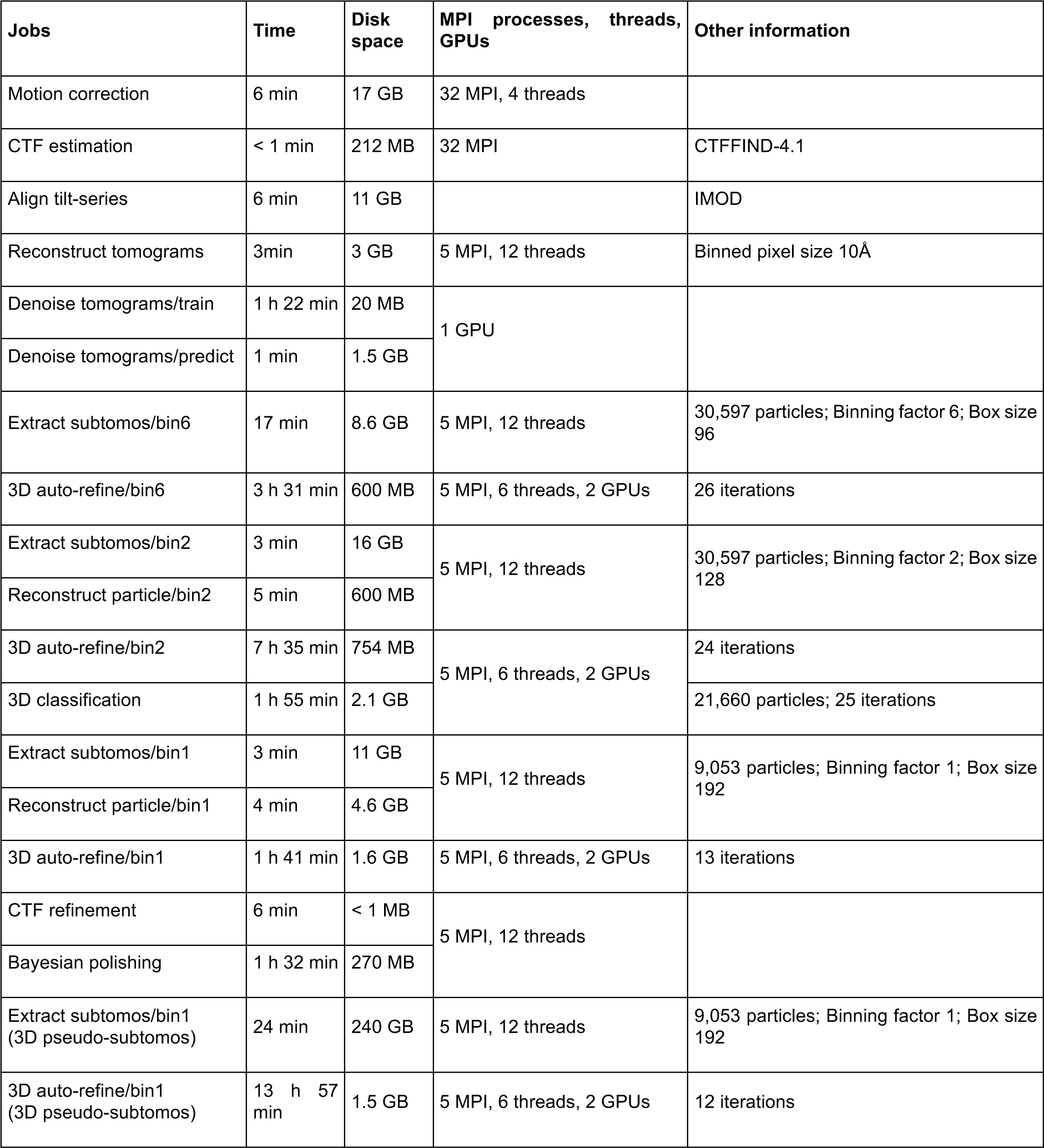
Computational costs. Each row represents a job described in the Results section, reporting its wall-clock time required for execution; how many MPI processes, threads and/or GPUs were used; and how much disk space the job required. All computations were performed on an (Ubuntu 22.04.3 LTS) Linux workstation with an AMD Ryzen Threadripper PRO 5975WX CPU with 32 cores and 64 threads, 512 GB RAM and two NVIDIA GeForce RTX 4090 GPUs, each with 24GB memory.

| Jobs | Time | Disk space | MPI processes, threads, GPUs | Other information |
| --- | --- | --- | --- | --- |
| Motion correction | 6 min | 17 GB | 32 MPI, 4 threads |  |
| CTF estimation | < 1 min | 212 MB | 32 MPI | CTFFIND-4.1 |
| Align tilt-series | 6 min | 11 GB |  | IMOD |
| Reconstruct tomograms | 3min | 3 GB | 5 MPI, 12 threads | Binned pixel size 10Å |
| Denoise tomograms/train | 1 h 22 min | 20 MB | 1 GPU |  |
| Denoise tomograms/predict | 1 min | 1.5 GB |  |  |
| Extract subtomos/bin6 | 17 min | 8.6 GB | 5 MPI, 12 threads | 30,597 particles; Binning factor 6; Box size 96 |
| 3D auto-refine/bin6 | 3 h 31 min | 600 MB | 5 MPI, 6 threads, 2 GPUs | 26 iterations |
| Extract subtomos/bin2 | 3 min | 16 GB | 5 MPI, 12 threads | 30,597 particles; Binning factor 2; Box size 128 |
| Reconstruct particle/bin2 | 5 min | 600 MB |  |  |
| 3D auto-refine/bin2 | 7 h 35 min | 754 MB | 5 MPI, 6 threads, 2 GPUs | 24 iterations |
| 3D classification | 1 h 55 min | 2.1 GB |  | 21,660 particles; 25 iterations |
| Extract subtomos/bin1 | 3 min | 11 GB | 5 MPI, 12 threads | 9,053 particles; Binning factor 1; Box size 192 |
| Reconstruct particle/bin1 | 4 min | 4.6 GB |  |  |
| 3D auto-refine/bin1 | 1 h 41 min | 1.6 GB | 5 MPI, 6 threads, 2 GPUs | 13 iterations |
| CTF refinement | 6 min | < 1 MB | 5 MPI, 12 threads |  |
| Bayesian polishing | 1 h 32 min | 270 MB |  |  |
| Extract subtomos/bin1 (3D pseudo-subtomos) | 24 min | 240 GB | 5 MPI, 12 threads | 9,053 particles; Binning factor 1; Box size 192 |
| 3D auto-refine/bin1 (3D pseudo-subtomos) | 13 h 57 min | 1.5 GB | 5 MPI, 6 threads, 2 GPUs | 12 iterations |

The results described above have been made available through Zenodo (DOI: 10.5281/zenodo.11068319) and form the basis for a detailed tutorial of the RELION-5 tomography pipeline, which can be found at http://relion.readthedocs.io.

## Discussion

Gaining structural insights from cryo-ET data is complex and remains an active area of research, with ongoing developments in CTF estimation ^39^, tilt series alignment ^8^, tomogram denoising ^21,23,24,45,52^, automated localisation of structures of interest ^30,53–62^ and approaches to deal with structural heterogeneity among extracted particles ^63–66^, amongst many others. As a result, cryo-ET image processing pipelines, including the one described here, will need to be continuously updated in the coming years. Probably the most urgent extensions to the pipeline in RELION-5 would include a more robust handling of reconstruction and defocus handedness from the provided metadata, more robust tools for automated tilt series alignment, and the incorporation of tools for automated tomogram segmentation and particle picking. Given the popularity of the combination of Warp/M ^50,67^ with sub-tomogram averaging in RELION-3 ^68^, updated inter-operability between Warp/M and RELION-5 is also desirable. Moreover, the use of napari as a platform for the development of these picking tools introduces complexity, especially for viewing data remotely. To avoid the maintenance burden of this complexity in RELION, the napari-based tools are intended to become standalone tools in the near future.

However, different research groups will require different tools to fulfil their specific image-processing needs, and it will remain difficult to include all of these in a single software pipeline. Recognising this, much of the work presented here was implemented as stand-alone python packages designed to maximise their reusability outside of the RELION framework. It would be helpful if the different groups that use and write cryo-ET software were to agree on a standardised description that allows passing metadata between different programs. In single-particle analysis, the exchange of metadata is often performed through files in the STAR format ^38^, with geometric definitions originally defined by Heymann et al ^69^ and then implemented in RELION ^70^. The explicit definition of metadata structures for tilt series, tomograms and extracted particles presented in this paper could fulfil a similar role for the cryo-ET field. A first step on this journey may be the adoption of RELION-5’s STAR files for cryo-ET data (Figures 2, 3 & 5 and Tables 1-3) in the upcoming cryo-ET pipeline of the Collaborative Computing Project for Electron cryo-Microscopy (CCP-EM) software ^37^. This new software will provide mechanisms for pipelining a wider range of cryo-ET programs than those available within RELION-5 (Tom Burnley, personal communication), thus further improving the accessibility for newcomers to this rapidly developing field.

## Conclusions

We present a pipeline for the analysis of cryo-ET data in RELION-5 that ranges from the import of unprocessed movies to automated atomic modelling in high-resolution sub-tomogram averaging maps. The explicit metadata definitions of tilt series, tomograms and extracted particles in RELION-5 may also serve wider efforts at standardisation and software inter-operability.

## Data accessibility

RELION is distributed under a GPLv2 open-source software license and can be downloaded for free from http://www.github.com/3dem/relion. All stand-alone python packages are distributed under a BSD-3 open-source software license. The data set described in the Results section can be downloaded from EMPIAR under accession number 10164, whereas the results themselves can be downloaded from Zenodo (DOI: 10.5281/zenodo.11068319).

## Author contributions

AB, BT, RW, EP, JZ, DK and SHWS wrote computer code; AB, BT, RW and SHWS performed experiments and analysed results. SHWS supervised the project. All authors contributed to writing the manuscript.

## Acknowledgements

We are grateful to Jake Grimmett, Toby Darling and Ivan Clayson for help with high-performance computing, and to Takanori Nakane for helpful discussions and RELION support. AB was a member of the group of David Barford (DB). This work was supported by the Medical Research Council as part of the United Kingdom Research and Innovation (MC_UP_A025_1013 to SHWS; MC_UP_1201/31 to TAMB and MC_UP_1201/6 to DB). AvK is supported by a Philip Leverhulme Prize to TAMB. RW is supported by The Pittsburgh Foundation & The Commonwealth of Pennsylvania Formula Fund to Zachary Freyberg; EP was supported by grants from the European Research Council (ERC-StG-2019 grant 852915) and the BBSRC (grant BB/T002670/1) to Giulia Zanetti. For the purpose of open access, the MRC Laboratory of Molecular Biology has applied a CC BY public copyright licence to any Author Accepted Manuscript version arising.

